# TOMM40 and TOMM22 of the Translocase Outer Mitochondrial Membrane Complex rescue statin-impaired mitochondrial dynamics, morphology, and mitophagy in skeletal myotubes

**DOI:** 10.1101/2023.06.24.546411

**Authors:** Neil V. Yang, Sean Rogers, Rachel Guerra, David J. Pagliarini, Elizabeth Theusch, Ronald M. Krauss

**Affiliations:** Department of Nutritional Sciences & Toxicology, University of California, Berkeley, CA, USA; Department of Pediatrics, University of California, San Francisco, CA, USA; Department of Medicine, University of California, San Francisco, CA, USA; Department of Cell Biology and Physiology, Washington University School of Medicine, St. Louis, MO, USA; Department of Biochemistry and Molecular BioPhysics, Washington University School of Medicine, St. Louis, MO, USA; Department of Genetics, Washington University School of Medicine, St. Louis, MO, USA

**Keywords:** Statin, mitochondrial dynamics, skeletal muscle, translocase of outer mitochondrial membrane, transmission electron microscopy

## Abstract

**Background:** Statins are the drugs most commonly used for lowering plasma low-density lipoprotein (LDL) cholesterol levels and reducing cardiovascular disease risk. Although generally well tolerated, statins can induce myopathy, a major cause of non-adherence to treatment. Impaired mitochondrial function has been implicated as a cause of statin-induced myopathy, but the underlying mechanism remains unclear. We have shown that simvastatin downregulates transcription of *TOMM40* and *TOMM22*, genes that encode major subunits of the translocase of outer mitochondrial membrane (TOM) complex which is responsible for importing nuclear-encoded proteins and maintaining mitochondrial function. We therefore investigated the role of *TOMM40* and *TOMM22* in mediating statin effects on mitochondrial function, dynamics, and mitophagy.

**Methods:** Cellular and biochemical assays and transmission electron microscopy were used to investigate effects of simvastatin and *TOMM40* and *TOMM22* expression on measures of mitochondrial function and dynamics in C2C12 and primary human skeletal cell myotubes.

**Results:** Knockdown of *TOMM40* and *TOMM22* in skeletal cell myotubes impaired mitochondrial oxidative function, increased production of mitochondrial superoxide, reduced mitochondrial cholesterol and CoQ levels, disrupted mitochondrial dynamics and morphology, and increased mitophagy, with similar effects resulting from simvastatin treatment. Overexpression of *TOMM40* and *TOMM22* in simvastatin-treated muscle cells rescued statin effects on mitochondrial dynamics, but not on mitochondrial function or cholesterol and CoQ levels. Moreover, overexpression of these genes resulted in an increase in number and density of cellular mitochondria.

**Conclusion:** These results confirm that TOMM40 and TOMM22 are central in regulating mitochondrial homeostasis and demonstrate that downregulation of these genes by statin treatment mediates disruption of mitochondrial dynamics, morphology, and mitophagy, effects that may contribute to statin-induced myopathy.

**GRAPHICAL ABSTRACT:** 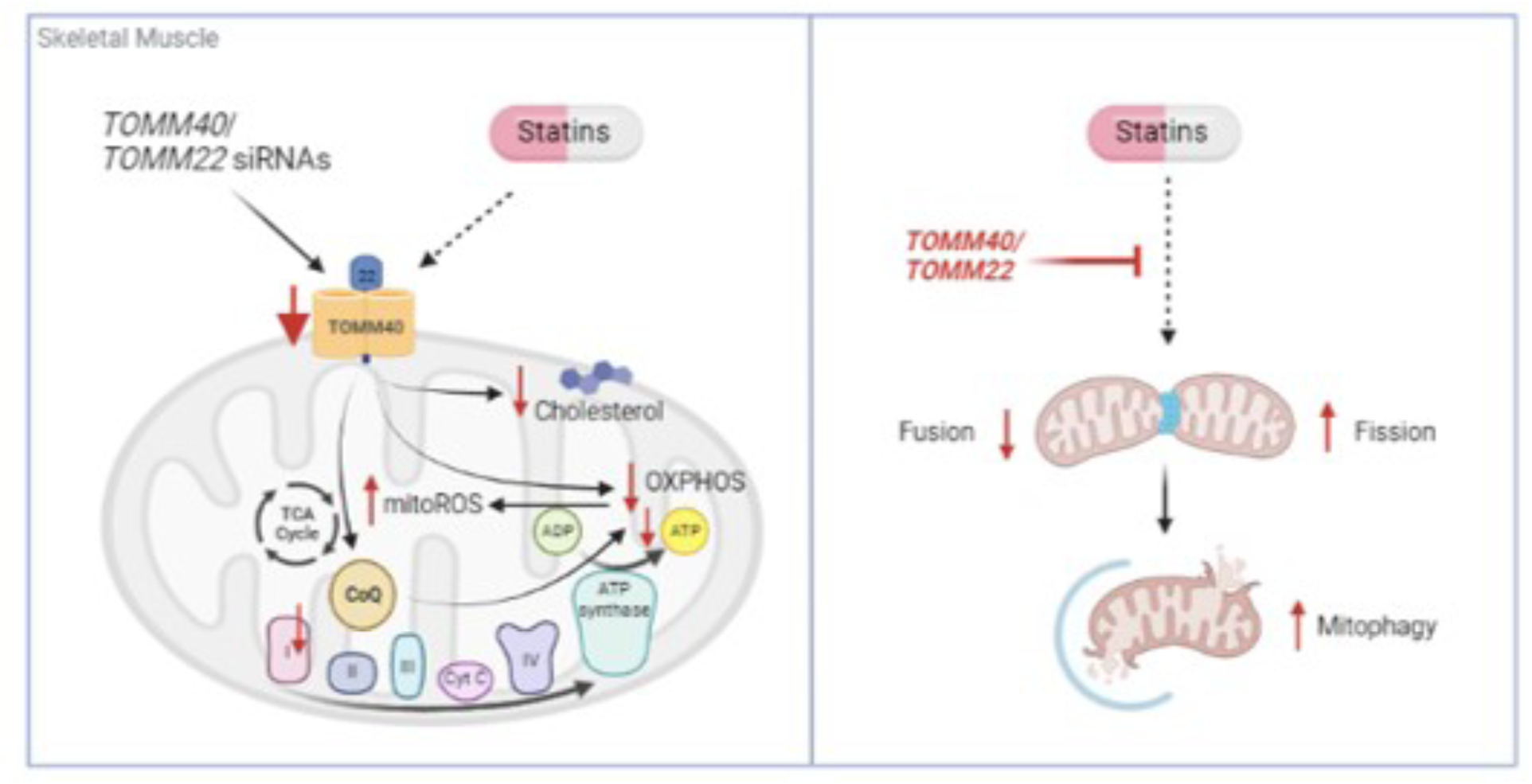

## INTRODUCTION

Statins, the most widely used class of drugs for reducing plasma LDL-cholesterol levels and cardiovascular disease risk, act by inhibiting 3-hydroxy-3-methyglutaryl coenzyme A reductase (HMGCR), the rate-limiting enzyme for cholesterol synthesis^1,2^. Although statins are highly effective and generally well-tolerated, they can have adverse side effects, the most common being statin-associated muscle symptoms (SAMS)^3,4^ ranging from myalgia and myositis to rhabdomyolysis^5^. Among the currently used statins, simvastatin has been associated with the greatest incidence of SAMS^6^.

Disruption of mitochondrial function has been proposed as a major mechanism contributing to SAMS^7^. Effects of statins on skeletal muscle mitochondria phenotypes have been demonstrated in cellular, animal, and clinical studies^8,9^. These include increased reactive oxygen species (ROS)^10^, decreased mitochondrial biogenesis^11^, altered protein prenylation, decreased intracellular ATP levels^12,13^, altered electron transport chain protein expression, increased fragmentation of mtDNA^14^, and elevated plasma creatine kinase (CK) levels^15^. Additionally, statin-induced inhibition of synthesis of coenzyme Q (CoQ_10_), an essential cofactor of the electron transport chain, has been proposed as a major contributing factor to skeletal muscle mitochondrial dysfunction^16^. Despite these findings, little is known of the molecular basis for these statin effects.

Recently, Grunwald et al. reported that simvastatin exposure of myotubes derived from primary human myoblasts resulted in a significant 50% reduction in expression of *TOMM40* and *TOMM22*^17^, two broadly expressed and highly conserved genes encoding the translocase of the outer mitochondrial membrane (TOM) complex^18^. We have observed a similar effect of simvastatin in a panel of human lymphoblastoid cell lines^19^. The mammalian TOM complex consists of 7 subunits that work together to recognize and import proteins from the cytoplasm into the mitochondrial interior to maintain mitochondrial function^20,21^ (**Fig. 1A**). Among these subunits, TOMM40 is the main channel-forming subunit that is stably associated with TOMM22, the central receptor of the complex^22^. Together, they are actively involved in protein translocation across the mitochondrial outer membrane^23^.

**Figure 1.**
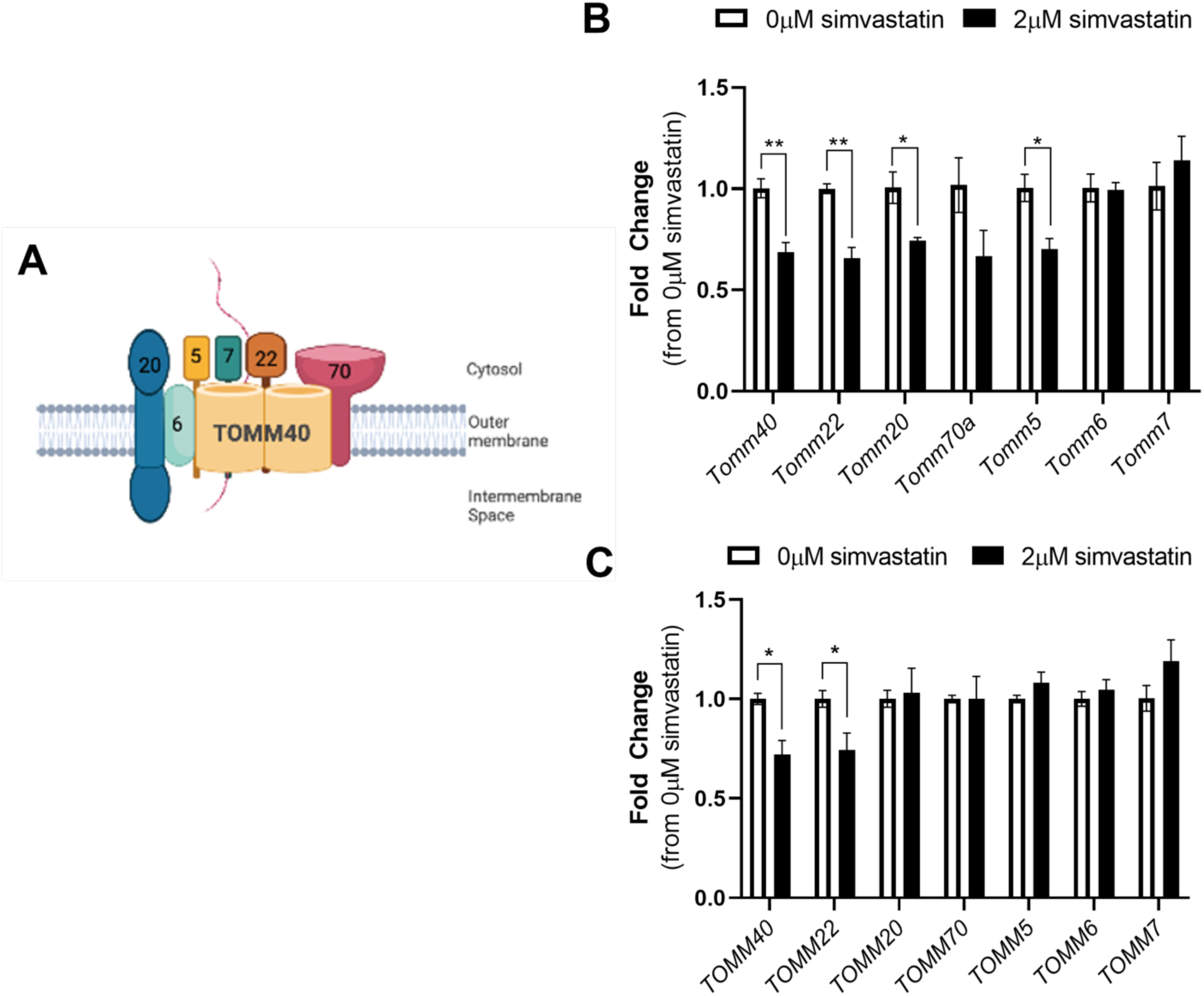
Simvastatin downregulates *TOMM40* and *TOMM22* in both C2C12 and hSkMC skeletal muscle. Differentiated C2C12 and primary hSkMC myotubes were treated with 2 µM simvastatin for 24 hrs. (A) A schematic diagram of the mammalian TOM complex, consisting of 7 subunits, located in the outer mitochondrial membrane. (B) Simvastatin treatment (2 µM) downregulates *Tomm40, Tomm22, Tomm20,* and *Tomm5* in differentiated C2C12 myotubes. (C) Simvastatin treatment (2 µM) downregulates *TOMM40 and TOMM22* in hSkMC myotubes. Numeric data represent mean ± SEM. \**p<0.05*, \*\**p<0.01*, \*\*\**p<0.001*, \*\*\*\**p<0.0001* vs. 0 µM simvastatin by Student’s t-test. (*n = 3* biological replicates).

In the present study, we have performed detailed mitochondrial phenotyping to compare the effects of both *TOMM40* and *TOMM22* knockdown and simvastatin treatment in two mammalian skeletal muscle cell models – mouse C2C12 cells, which are known to display cellular phenotypes with statin treatment similar to those seen with statin-induced myotoxicity in humans^24^ and primary human skeletal muscle cells (hSkMC). We also tested the role of *TOMM40* and *TOMM22* downregulation in mediating statin’s mitochondrial effects by determining whether these effects were rescued by overexpressing these genes in statin treated skeletal muscle cells.

## METHODS

### Cell Culture

C2C12 murine myoblasts and primary hSkMCs were purchased from American Type Culture Collection (ATCC; CRL-1772 & PCS-950-010). C2C12 cells were cultured in DMEM containing 4.5 g/L glucose and L-glutamine (Gibco) supplemented with 10% fetal bovine serum (FBS; Thermo Fisher Scientific) and penicillin-streptomycin (Gibco) at 37°C and 5% CO_2_. After passaging cells, C2C12 myoblasts were differentiated into myotubes by replacing media with DMEM containing 2% horse serum (Gibco). Fresh medium was replaced every 2 days and cells were incubated for 5-7 days to completely differentiate into myotubes before experimentation. For hSkMCs, cells were cultured in mesenchymal stem cell basal medium (PCS-500-030, ATCC) supplemented with L-glutamine, 5 ng/mL rh EGF, 10 µM dexamethasone, 5 ng/mL rh FGF-b, 25 µg/mL rh insulin, 4% FBS (PCS-950-040, ATCC) and penicillin-streptomycin (Gibco). 24-96 hrs after passaging, hSkMC cells were differentiated into myotubes using skeletal muscle differentiation tool (PCS-950-050, ATCC) for 2 days before experimentation. Cells were routinely tested for mycoplasma using MycoAlert™ PLUS mycoplasma detection kit (Lonza) and only mycoplasma negative cells were used.

### siRNA Reverse Transfections

To achieve knock-down (KD) of *TOMM22* and *TOMM40*, C2C12 and hSkMC cells were seeded at 100,000 cells per well in 6-well plates. Upon seeding, cells were reverse transfected with 10 µM of non-targeting control (NTC), *TOMM22* and/or *TOMM40*-targeted siRNAs using Lipofectamine RNAiMax transfection reagent (Thermo Fisher Scientific) and Opti-MEM 1 (Gibco) for 48hrs. All siRNAs were purchased from Thermo Fisher Scientific – human *TOMM22* siRNA (s32549); human *TOMM40* siRNA (s20449); mouse *Tomm22* siRNA (s104588); mouse *Tomm40* siRNA (s79125); Silencer Select Negative Control siRNA (s79125). For C2C12, cells were transfected twice, at day 0 (myoblasts) and day 3 (myotubes) in differentiation media and then harvested after day 5 for experimentation.

### Overexpression Plasmids

Human and mouse pCMV-EGFP expressing-*TOMM40/Tomm40* (human: NM_001128916.2, mouse: NM_016871.2), *TOMM22/Tomm22* (human: NM_020243.5, mouse: NM_172609.3), *Tomm20* (mouse: NM_024214.2), and empty vector (EV; ORF_stuffer) plasmids stored in bacterial glycerol stocks were purchased from VectorBuilder Inc. Expression plasmids were cultured on Luria-Bertani (LB) Agar plates containing ampicillin at 37°C. Single colonies were selected and grown separately in LB broth at 37°C with continuous shaking (225 rpm) overnight. DNA plasmids were then purified and extracted using the ZymoPURE II Plasmid Midiprep Kit (Zymogen) according to manufacturer’s protocol. For overexpression studies, both C2C12 and hSkMC cells were first differentiated into myotubes using respective differentiation media. After differentiation, cells were transiently transfected with purified *TOMM40/Tomm40*, *TOMM22/Tomm22,* and *Tomm20* expression plasmids, singly and in combination, or matched empty vector, using Lipofectamine 3000 transfection reagent (Thermo Fisher Scientific). In parallel, 2 µM simvastatin was added to the cell media and cells were collected after 24-48 hrs. Simvastatin was obtained as a gift from Merck and activated as previously described^25^.

### Mitochondrial Respiration Measurements

*In vitro* oxygen consumption rate was measured in fully differentiated C2C12 and hSkMC skeletal myotubes with the Agilent Seahorse XFe96 Extracellular Flux Analyzer. Cells were seeded at 1,000 per well in 96-well plates, with XF assay medium (Agilent) supplemented with 2 mM sodium pyruvate (Gibco), 2 mM GlutaMAX™ (Gibco), and 10 mM glucose (Sigma), at pH 7.4. During experimentation, 1.5 µM oligomycin, 2 µM FCCP, and 2 µM Antimycin A + Rotenone (Seahorse XF Cell Mito Stress Test Kit, Agilent) was added sequentially via injection ports, to calculate basal and maximum respiration and ATP production. Oxygen consumption rate (OCR) values were presented with non-mitochondrial oxygen consumption deducted and normalized to total protein concentration per well using Bradford assay.

### Fluorescence Quantification

To detect mitochondria superoxide production in C2C12 cells, myotubes were incubated with 5 µM MitoSOX™ Red (Molecular Probes) for 20 min at 37°C. Cells were then rinsed twice in pre-warmed 1X phosphate-buffered saline (PBS) which was then replaced with phenol red-free DMEM (21063029; Gibco) supplemented with glucose and sodium pyruvate. Fluorescence was detected and quantified at an excitation/emission of 396/610 nm using an Agilent BioTek™ microplate fluorescence spectroscopy reader and BD LSRFortessa™ Cell Analyzer flow cytometer following the protocol of Kauffman et al^26^. To quantify mitochondrial density, MitoTracker™ Deep Red FM (100 nM) was added to C2C12 cells for 30 min at 37°C, washed twice with 1X PBS and replaced with phenol red-free DMEM supplemented with glucose and sodium pyruvate (Gibco). MitoTracker fluorescence was quantified at an excitation/emission of 644/665 nm. All absorbance readings were normalized to total protein concentration by Bradford assay.

### Isolation of Mitochondria by Subcellular Fractionation

Mitochondria were isolated from C2C12 and hSkMC cells according to the method of Wettmarshausen and Perocchi^27^. Cells were rinsed in 1X PBS twice, dislodged with 0.25% Trypsin-EDTA (Gibco), washed again in 1X PBS and centrifuged at 600 x g for 5 min at 4°C. Pelleted cells were resuspended in MSHE + BSA buffer (210 mM mannitol, 70 mM sucrose, 5 mM HEPES, 1 mM EGTA, and 0.5% BSA, at 7.2 pH). Samples were transferred to a small glass dounce and homogenized. The homogenate was centrifuged at 600 x g for 10 min at 4°C, and the supernatant was extracted and centrifuged at 8,000 x g for 10 min at 4°C. The isolated pellet containing the purified crude mitochondria was dried down by nitrogen gas and snap frozen in liquid nitrogen for storage.

### Lipid Extraction for LC-MS/MS

C2C12 whole cell lysates and isolated mitochondria pellets were resuspended in 100 µL of 150 mM KCl. Ten percent of the cell suspension was removed from each tube and placed into a new tube. The extra 10% was later used in a BCA assay to measure relative protein content in each sample. Protein content derived from the BCA assay was used to normalize CoQ measurements.

The remaining 90% of the cell suspension was mixed with glass beads and 600 µL of cold methanol containing 0.25 µM CoQ_6_ (CoQ_6_ is used to normalize for total CoQ extracted). Cell suspensions were subjected to lysis on a vortex genie at 4°C for 10 min. Afterward, 400 uL of cold petroleum ether was added to each tube and vortexing was repeated for 3 min. To separate the petroleum ether and methanol phases, the tubes were centrifuged at 1,100 x g for 3 min and the top (petroleum ether) phase was collected into a new tube (Tube B). Again, 400 µL of petroleum ether was added to each tube containing methanol, and the vortexing/centrifuge steps were repeated. The final top layer was collected and added to Tube B. Petroleum ether was dried under a stream of argon gas, and dried lipids were resuspended in 50 µL of mobile phase (78:20:2 methanol:isopropanol:ammonium acetate).

### Measurement of CoQ_9_ by LC-MS/MS Lipidomics

#### LC-MS/MS Lipidomics Data Acquisition

A Vanquish Horizon UHPLC system (Thermo Scientific) connected to an Exploris 240 Orbitrap mass spectrometer (Thermo Scientific) was used for targeted LC-MS analysis. A Waters Acquity CSH C18 column (100 mm × 2.1 mm, 1.7 μm) was held at 35°C with the flow rate of 0.3 mL/min for lipid separation. A Vanquish binary pump system was employed to deliver mobile phase A consisting of 5 mM ammonium acetate in ACN/H2O558(70/30, v/v) containing 125 μL/L acetic acid, and mobile phase B consisting of 5 mM ammonium acetate in IPA/ACN (90/10, v/v) containing 125 μL/L acetic acid. The gradient was set as follows: B was at 2% for 2 min and increased to 30% over the next 3 min, then further ramped up to 50% within 1 min and to 85% over the next 14 min, and then raised to 99% over 1 min and held for 4 min, before being re-equilibrated for 5 min at 2% B. Samples were ionized by a heated ESI source with a vaporizer temperature of 350°C. Sheath gas was set to 50 units, auxiliary gas was set to 8 units, sweep gas was set to 1 unit. The ion transfer tube temperature was kept at 325°C with 70% RF lens. Spray voltage was set to 3,500 V for positive mode. The targeted acquisition was performed with tMS2(targeted MS2) mode: tMS2 mode was for measuring CoQ_6_(m/z 591.4408, internal standard) and CoQ_9_(m/z 795.6286) in positive polarity at the resolution of 15,000, isolation window of 2m/z, normalized HCD collision energy of either 40% or stepped HCD energies of 30% and 50%, standard AGC target and auto maximum ion injection time.

#### Data Analysis

Targeted quantitative analysis of all acquired compounds was processed using TraceFinder 5.1 (Thermo Scientific) with the mass accuracy of 5 ppm. The result of peak integration was manually examined.

### Lipid Extraction and Intracellular Cholesterol Quantification

Cholesterol was extracted from cells with hexane-isopropanol (3:2, v/v), dried under nitrogen gas and reconstituted with buffer (0.5 M potassium phosphate, pH 7.4, 0.25 M NaCl, 25 mM cholic acid, 0.5% Triton X-100). Intracellular cholesterol levels were then quantified with the Amplex Red Cholesterol Assay Kit (Life Technologies) according to the manufacturer’s protocol.

### Sample Preparation for Electron Microscopy

C2C12 cells were grown on MatTek glass bottom dishes (P35G-1.5-14-C, MatTek) and fixed in 2% glutaraldehyde + 2% paraformaldehyde solution (prepared by Electron Microscopy Lab, UC Berkeley) for 24 hrs. After fixation, cells were washed 3-times for 5 min in 0.1 M sodium cacodylate buffer, pH 7.4. Samples were then post-fixed in 1% osmium tetroxide + 1.6% potassium ferricyanide (KFECn) in 0.1 M sodium cacodylate buffer for 30 min, before undergoing 3 washes at 15 min each. Cells were then dehydrated in a serial diluted ethanol solution of 30, 50, 70, 90, and 100%, for 10 min each. Samples were infiltrated with 50% Epon-Araldite resin (containing benzyldimethylamine (BDMA) accelerator), followed by 100% resin for 1 hr each. Excess resin was removed from the MatTek dishes containing cells and polymerized at 60°C for 48 hrs.

### Transmission Electron Microscopy

Using a dissecting blade, cells embedded in resin were removed from MatTek dishes and mounted on resin-embedded blocks for sectioning. Serial sections of 70-150 nm thickness were cut on a Reichert-Jung Ultracut E microtome and set on 1 x 2-mm slot grids covered with 0.6% Formvar film. Sections were then post-stained with 1% aqueous uranyl acetate for 7 min and lead citrate for 4 min^28^. Images of cell samples were taken on an FEI Tecnai 12 transmission electron microscope equipped with a 2k x 2k CCD camera with a 40 Megapixel/sec readout mode. Images were analyzed using ImageJ software according to the method by Lam et al^29^.

### Immunoblotting

Cells were washed with PBS and lysed in M Cellytic Lysis Buffer containing 1% protease inhibitor (Halt ™ Protease Inhibitor Cocktail; Thermo Scientific) for 15 min with gentle vortexing. The cell lysate was centrifuged at 14,000 x g for 15 min, the supernatant was collected, and the protein concentration was measured by Bradford assay. Proteins were separated on a 4-20% Tris-polyacrylamide gradient gel (Bio-Rad) and transferred onto a nitrocellulose membrane using the iBlot™ 2 Gel Transfer Device (Thermo Fisher Scientific). Membranes were blocked in Tris-buffered saline with 0.1% tween (TBST) + 5% milk for 2 hrs to minimize non-specific antibody binding. Membranes were then incubated with primary antibodies diluted in TBST overnight on a rotating platform at 4°C. After washing in TBST, membranes were incubated with secondary antibodies for 30 min before a last series of washes. SuperSignal™ West Pico PLUS Chemmiluminescent Substrate (Thermo Fisher Scientific) was added to the membrane to visualize proteins^30^.

### RT-qPCR and mtDNA Copy Number

RNA was extracted from cells using RNeasy Mini Qiacube Kit (Qiagen) with the Qiacube Connect (Qiagen) according to manufacturer’s protocol. cDNA synthesis from total RNA was performed using High Capacity cDNA Reverse Transcription Kits (Applied Biosystems). Primers obtained from Elim Biopharmaceuticals were run with SYBR™ Green qPCR Master Mix (Thermo Fisher Scientific) on an ABI PRISM 7900 Sequence Detection System to quantify mRNA transcript levels. RT-qPCR primers used in this study are listed in **Table S1**. The mean value of triplicates for each sample was normalized to GAPDH as the housekeeping gene.

Total DNA was isolated from C2C12 cells using DNeasy Blood and Tissue Kit (Qiagen). qPCR was performed with SYBR™ Green qPCR Master Mix (Thermo Fisher Scientific) on an ABI PRISM 7900 Sequence Detection System according to the protocol outlined by Quiros et al^31^. Primers for mouse mtDNA (mMitoF1: *5’-CTAGAAACCCCGAAACCAAA-3’*, mMitoR1: *5’- CCAGCTATCACCAAGCTCGT-3’*) and mouse B2M (mB2MF1: *5’- ATGGGAAGCCGAACATACTG-3’*, mB2MR1: *5’CAGTCTCAGTGGGGGTGAAT-3’*) were used to amplify mtDNA and nuclear DNA, respectively. mtDNA copy number was determined by normalizing mtDNA to nuclear DNA. The delta delta Ct method was used to calculate fold change in mtDNA copy number.

### Statistical Analysis

All data are presented as the mean ± standard error of mean (SEM). *N*-values in the figures refer to biological replicates and at least 3 replicates were conducted per condition and experiment. *P*-values were calculated using Student’s t-tests for two groups. To compare more than two groups, one-way analysis of variance (ANOVA) or Welch and Brown-Forsythe ANOVA with Tukey’s post hoc test were used. Analyses were performed using GraphPad Prism 9 software (GraphPad Software, Inc.). *P<0.05* were considered statistically significant.

## RESULTS

### Simvastatin downregulates key subunits of the TOM complex in mammalian skeletal muscle cells

We first aimed to confirm previous findings in primary human myotubes^23^ that expression of *TOMM40* and *TOMM22,* subunits of the TOM complex, are downregulated by simvastatin exposure. Differentiated C2C12 and primary hSkMC myotubes were treated with 2 µM of simvastatin for 24 hrs. This dose was chosen based on simvastatin dose response experiments that showed significant induction of *Hmgcr* and *Ldlr* mRNA at 2 µM in C2C12 cells (**Fig. S1**). Additionally, we performed a simvastatin dose response experiment in C2C12 cells to assess apoptosis using EarlyTox Caspase-3/7 and found no significant increase with simvastatin 2 µM vs. baseline **(Fig. S2**). In both cell types, mRNA transcript levels of *TOMM40* and *TOMM22* were significantly reduced by exposure to 2 µM simvastatin as assessed by qRT-PCR **(Fig. 1B, C).** Transcript levels of two other subunits of the TOM complex, *Tomm20* and *Tomm5*, were also significantly reduced by simvastatin in C2C12 myotubes **(Fig. 1B).** These results confirm and extend evidence that simvastatin downregulates expression of the major subunits of the TOM complex.

### *TOMM40* and *TOMM22* knockdown impair mitochondrial function in skeletal myotubes

We next sought to assess the potential role of *TOMM40* and *TOMM22* downregulation by statins in mediating mitochondrial dysfunction by studying the effects of *TOMM40* and *TOMM22* knockdown (KD) in C2C12 and hSkMC myotubes. KD of *Tomm40* and *Tomm22* singly and in combination was performed by a two-step siRNA transfection (**Fig. 2A**), resulting in greater than ∼80% knockdown efficiency (**Fig. 2B**). In both C2C12 and hSkMC myotubes, basal and maximal oxygen consumption rate (OCR), as well as ATP production, were significantly reduced by each condition compared with a non-targeting control (NTC), indicating impaired electron transport chain and mitochondrial function (**Fig. 2C-F**).

**Figure 2.**
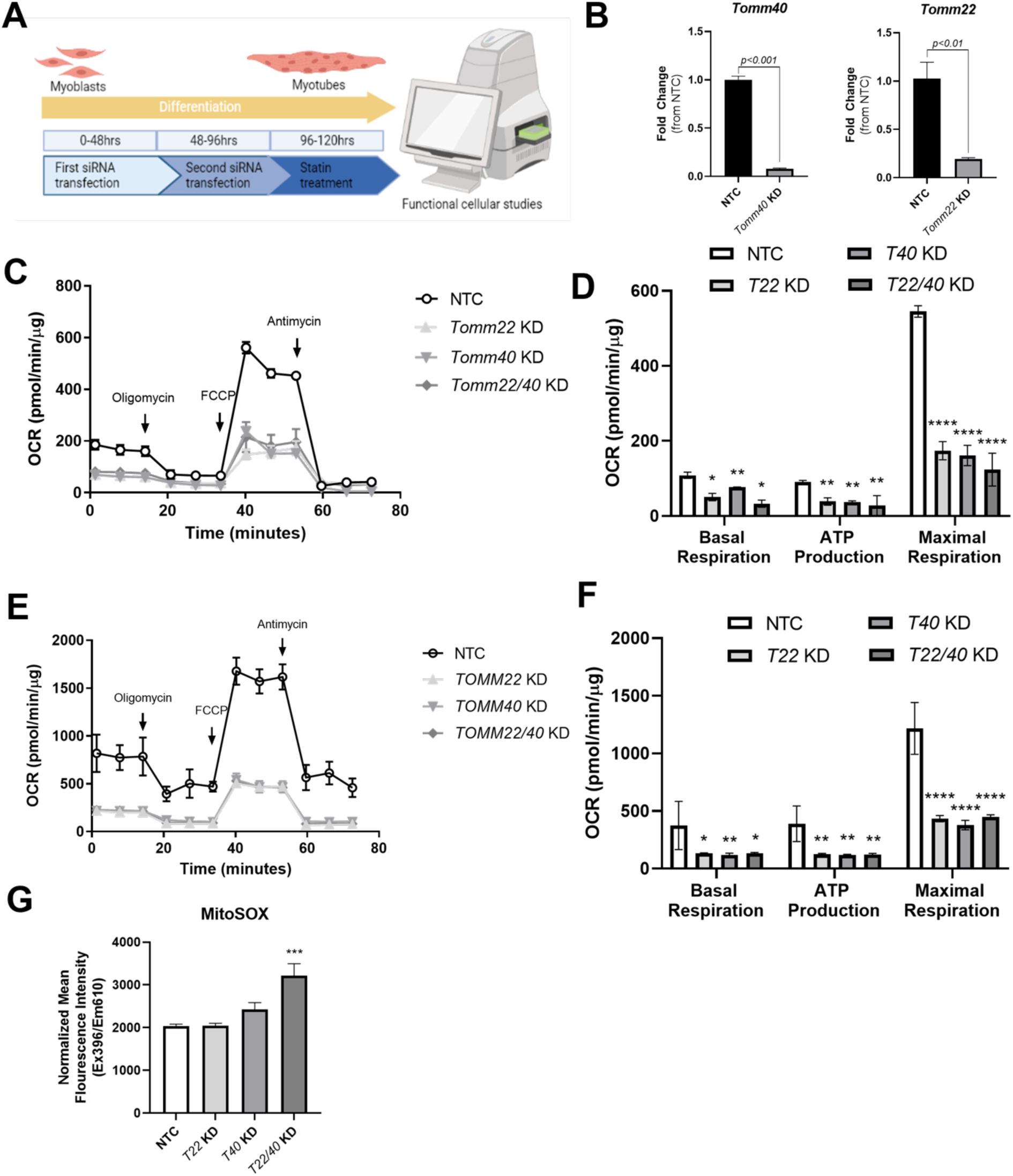
*TOMM40* and *TOMM22* knock-down decrease mitochondrial oxygen consumption rate and promote mitochondrial superoxide production. (A) Schematic illustration of the two-step transfection and differentiation experiment in C2C12 cells, *in vitro*. (B) Confirmation of *Tomm40* and *Tomm22* KD in C2C12 myotubes by qPCR. (C) Oxygen consumption rates of C2C12 myotubes transfected with Tomm40 and Tomm22 siRNAs vs. NTC were quantified using the Seahorse 96e Extracellular Flux Analyzer. With the addition of oligomycin, FCCP, and Antimycin A + Rotenone, basal respiration, ATP production, and maximal respiration were quantified. (D-F) The same experiment done in C2C12 cells (C-D) was conducted in primary hSkMC cells. (G) Mitochondrial superoxide production with *Tomm40* and *Tomm22* KD in C2C12 cells was quantified by MitoSOX fluorescence probe. All values were normalized to protein concentration by Bradford assay. All graphical and numeric data represent mean ± SEM. \**p<0.05*, \*\**p<0.01*, \*\*\**p<0.001*, \*\*\*\**p<0.0001* vs. NTC by Student’s t-test. (*n = 10-12* biological replicates).

To confirm a decrease in mitochondrial respiration due to suppression of *TOMM40* and *TOMM22* gene expression, we measured mitochondrial superoxide (mitochondrial reactive oxygen species, a.k.a. mitoROS) production using a fluorescence indicator (MitoSOX™) in C2C12 myotubes. This showed increased mitochondrial superoxide production in C2C12 cells transfected with both *Tomm40* and *Tomm22* siRNAs, compared to NTC (**Fig. 2G**). Though there was an increased trend observed in cells transfected with *Tomm40* siRNA individually (*p*=0.171), no significant differences were observed in *Tomm22* KD cells alone. These results demonstrate that KD of either *TOMM40* or *TOMM22* in skeletal myotubes impairs mitochondrial respiration and that their combined KD promotes the generation of mitoROS.

### *Tomm40* and *Tomm22* KD reduce cholesterol and CoQ levels in mitochondria of C2C12 myotubes *in vitro*

Since mitochondria require cholesterol for maintenance of membrane integrity and proper respiratory function^32^, we first tested whether suppression of *Tomm40* and *Tomm22* expression impairs mitochondrial cholesterol levels, and second if *Tomm40* and *Tomm22*-induced changes on mitochondrial cholesterol content contributes to the impaired mitochondrial respiration observed. Cholesterol content was analyzed by colorimetric assay in mitochondria isolated from C2C12 myotubes by subcellular fractionation **(Fig. 3A**). Consistent with our hypothesis, a reduction in mitochondrial cholesterol content was observed in both *Tomm40* and *Tomm22* siRNA-transfected cells, singly and in combination, compared to NTC (**Fig. 3B**). To assess whether reduced cholesterol alone is responsible for the disruption of mitochondrial function by *Tomm40* and *Tomm22* KD, we performed a cholesterol addback experiment. The addition of LDL isolated from human plasma rescued mitochondrial cholesterol content of *Tomm22* KD but not *Tomm40* KD myotubes (**Fig. 3C, D**). However, cholesterol repletion in *Tomm*22 KD myotubes did not restore reduced mitochondrial ATP production and basal respiration (**Fig. 3E, F**).

**Figure 3.**
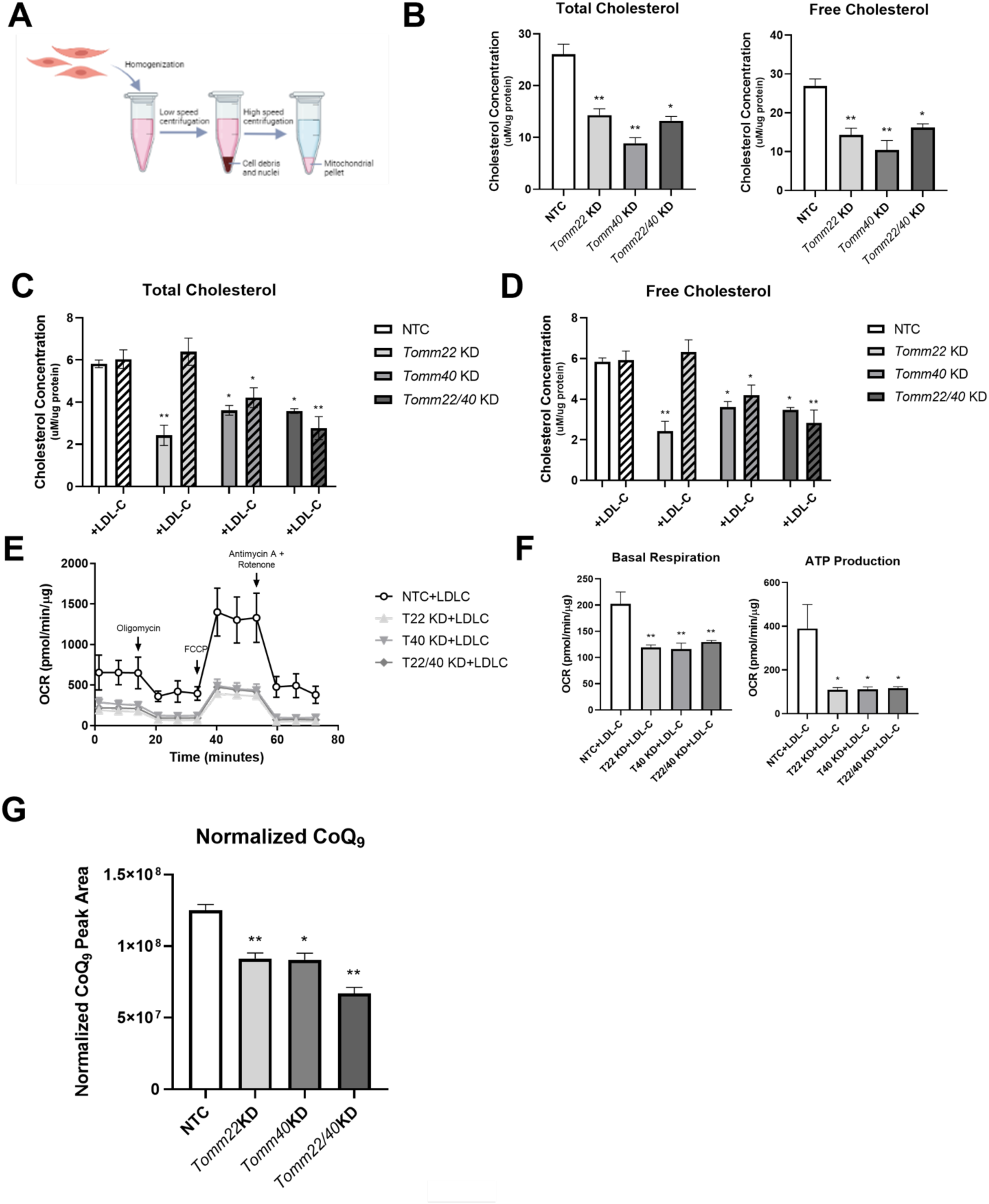
*Tomm40* and *Tomm22* regulate mitochondrial cholesterol content and CoQ levels in C2C12 myotubes. (A) Subcellular fractionation was performed to isolate crude mitochondria from whole cells. (B) Total and free cholesterol levels in mitochondria isolated from C2C12 myotubes transfected with NTC, *Tomm40*, and *Tomm22* siRNAs, singly and in combination. (C-D) In a separate experiment, after cells were transfected with siRNAs, 50 µg/mL LDL-C was added to cell media for 24 hrs. Total and free cholesterol were quantified in the mitochondria using Amplex Red Cholesterol Assay. (E) OCR levels were assessed in C2C12 cells after the LDL-C addback and (F) basal respiration and ATP production were quantified using Wave software (*n = 10-12* biological replicates). (G) Total CoQ from isolated mitochondria of NTC, *Tomm40*, *Tomm22*, and *Tomm22/40* KD C2C12 cells were quantified by LC-MS/MS. All values were normalized to protein concentration, measured by BCA. All graphical and numeric data represent mean ± SEM \**p<0.05*, \*\**p<0.01*, \*\*\**p<0.001*, \*\*\*\**p<0.0001* vs. NTC (without LDL-C addback) by Welch and Brown-Forsythe ANOVA or Student’s t-test. (*n = 3* biological replicates).

Given that CoQ, a product of the CoQ biosynthesis pathway, plays a central role in the mitochondrial electron transport chain, we next tested the effects of *Tomm40* and *Tomm22* KD on mitochondrial CoQ levels. Since mice unlike humans predominantly synthesize CoQ_9_ (9 prenyl units), we analyzed CoQ_9_ levels in our murine C2C12 cell model^33^. While *Tomm40* and *Tomm22* KD, singly and in combination, resulted in no differences in levels of CoQ (**Fig. S3**) in whole cell lysates, there were significant reductions in CoQ_9_ in isolated mitochondria (**Fig. 3G**). Together, these results indicate that TOMM40 and TOMM22 of the TOM complex may impact mitochondrial function in skeletal myotubes at least in part by its effect on the CoQ biosynthesis pathway.

### *TOMM40* and *TOMM22* KD promote mitochondrial fission and upregulate mitophagy in response to mitochondrial damage

Having observed mitochondrial dysfunction with the suppression of *TOMM40* and *TOMM22*, we next investigated mitochondrial dynamics, a key process that regulates cellular and mitochondrial metabolism^34^. By transmission electron microscopy (TEM) of C2C12 myotubes transfected with *Tomm22* and *Tomm40* siRNAs compared to NTC, we observed an increase in constriction events within individual mitochondria, indicative of increased fission events (**Fig. 4A**), along with a significant decrease in average mitochondrial length (**Fig. 4B**) and no change in average width (**Fig. 4C**). This change in mitochondrial morphology led us to hypothesize that *Tomm40* and *Tomm22* KD affect mitochondrial dynamics in skeletal muscle. By qPCR in *TOMM22* and *TOMM40* KD C2C12 and hSkMC cells we observed upregulated expression of *FIS1* and *DNM1L/DRP1*, genes that encode markers of mitochondrial fission (**Fig. 4D, E**). This was accompanied by a decrease in gene expression of the mitochondrial fusion markers *MFN2* and *OPA1*. With a shift towards mitochondrial fission in *TOMM22* and *TOMM40* KD skeletal muscle cells, we observed an increase in mitochondrial density (**Fig. 4F**) and mtDNA copy number (**Fig. 4G**), which together indicate increased mitochondrial fragmentation and support the upregulated mitochondrial fission observed.

**Figure 4.**
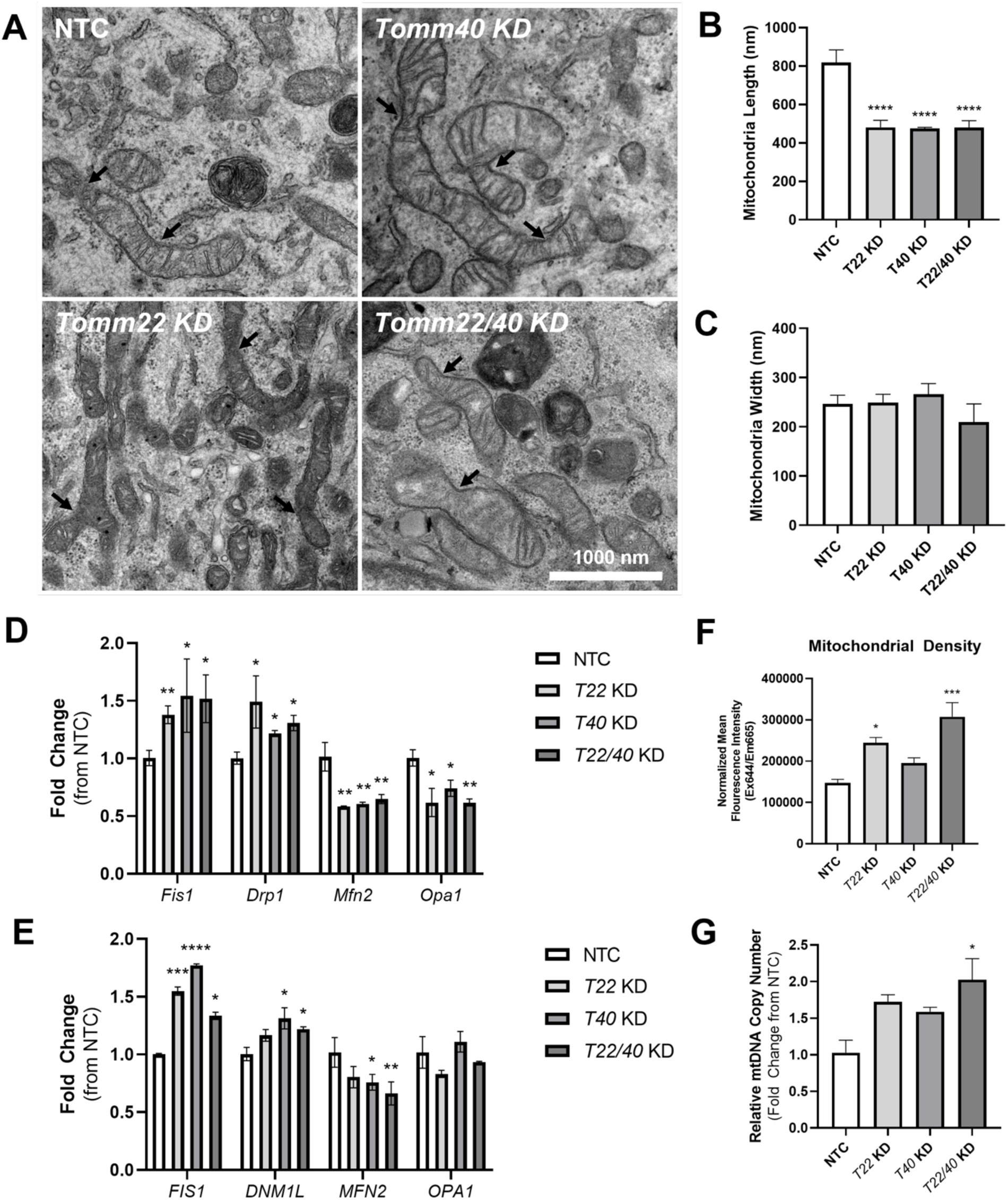
*TOMM40* and *TOMM22* knockdown, singly and in combination, impair mitochondrial dynamics in skeletal myotubes. (A) TEM micrographs of NTC, Tomm40, Tomm22, and Tomm22/40 KD in C2C12 cells. Arrowheads indicate mitochondrial fission events. Analysis of mitochondrial morphology using ImageJ software: (B) average mitochondrial length (nm) and (C) average mitochondrial width (nm). *n = 10-12* cells. (D) Mitochondrial fission (*Fis1/FIS1, Drp1/DNM1L*) and fusion (*Mfn2/MFN2, Opa1/OPA1*) markers were quantified by qPCR in NTC, Tomm22, Tomm40, and Tomm22/40 KD C2C12 cells. (E) The experiment was conducted as in (D) but with hSkMC cells. (*n=3* biological replicates) (F) Mitochondrial density was quantified using MitoTracker™ Deep Red FM fluorescence probe to measure average fluorescence intensity and normalized to protein concentration by Bradford assay (n=10-12 biological replicates). (G) Relative mtDNA copy number levels were quantified and normalized to *B2m* transcript levels (nuclear DNA) by qPCR in siRNA transfected C2C12 cells (*n = 3* biological replicates). All graphical and numeric data represent mean ± SEM. \**p<0.05*, \*\**p<0.01*, \*\*\**p<0.001*, \*\*\*\**p<0.0001* vs. NTC by Student’s t-test.

We further used TEM for analyzing mitochondrial morphology to assess mitochondrial damage (**Fig. 5A**). KD of *Tomm40* and *Tomm22* resulted in a reduced percentage of type 1 (healthy) mitochondria, and an increase in both type 2 and 3 (damaged and ruptured) mitochondria (**Fig. 5B, C**)^35^. Collectively, these results indicate that while there is an increase in new mitochondria created from fission events in knock-down cells (as confirmed in **Fig. 4F, G**), the majority are damaged. Furthermore, we noticed a significant increase in percentage of mitophagosomes in *Tomm40* and *Tomm22* KD C2C12 cells (**Fig. 5D**). This observation was confirmed by an increase in gene and protein expression of the mitophagy markers *PINK1* and *PRKN* in hSkMCs (**Fig. 5E, F**). Accordingly, these results support a compensatory mechanism due to the mitochondrial damage induced by *TOMM40* and *TOMM22* KD in which mitochondrial fission is upregulated leading to increased mitophagy that removes damaged mitochondria and maintains mitochondrial homeostasis.

**Figure 5.**
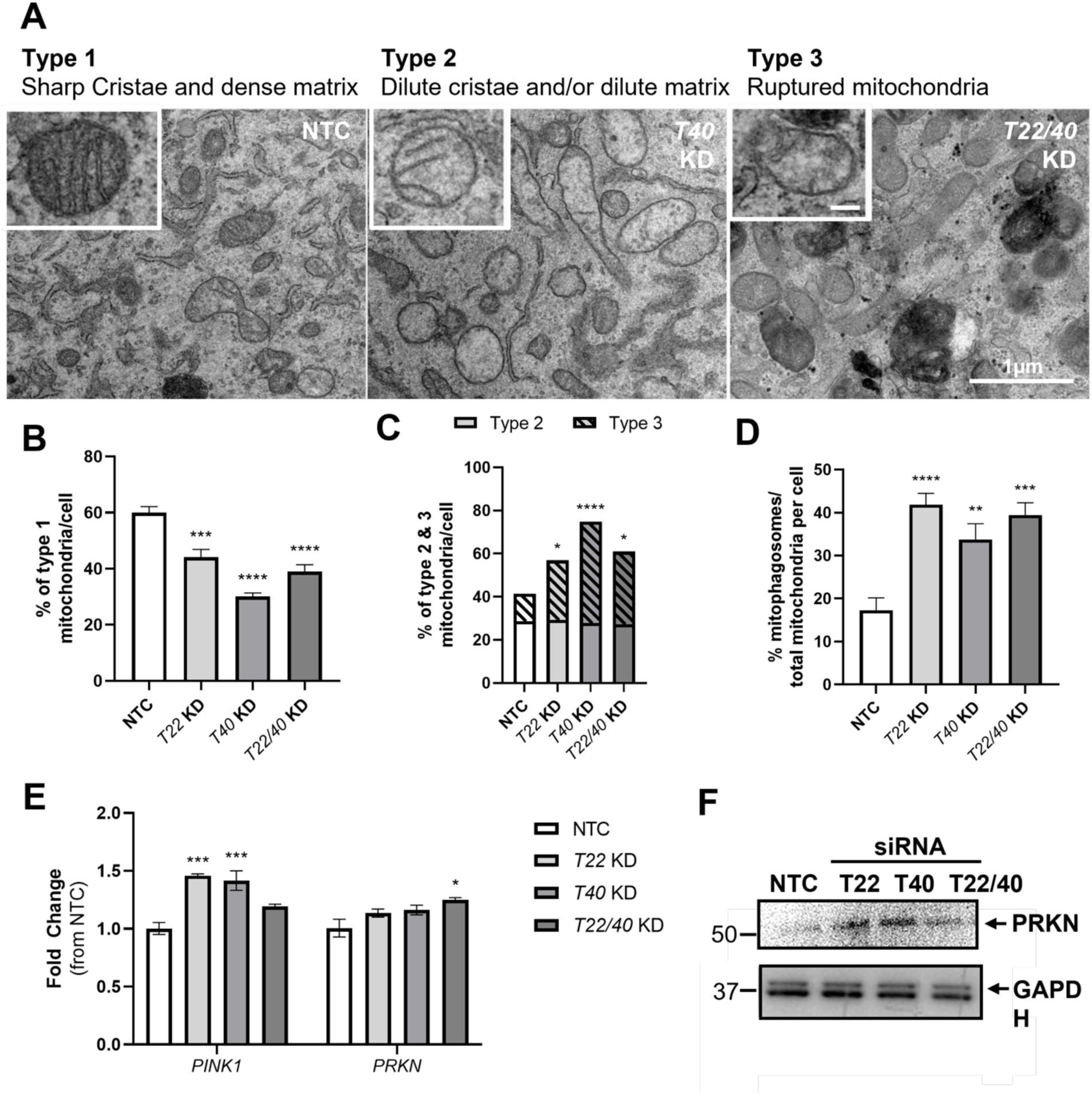
*TOMM40* and *TOMM22* knock-down, singly and in combination, promote mitochondrial damage and mitophagy in skeletal myotubes. (A) TEM micrographs of NTC, Tomm40, and Tomm22/40 KD C2C12 myotubes representing Type 1 (healthy), 2 (unhealthy), and 3 (damaged/ruptured) mitochondria, respectively (inset scale bar = 200nm). (B) Analysis of mitochondrial morphology and damage in NTC vs. KD C2C12 cells from (A). Bar graph represents percent of cells exhibiting type 1 mitochondria. (C) Bar graph represents percent of cells exhibiting type 2 and 3 mitochondria (*n = 10-12* cells). (D) Percent of mitophagosomes per total number of mitochondria per cell, identified from TEM images (*n = 10-12* cells). (E) mRNA transcript levels of *PINK1* and *PRKN* (mitophagy), were quantified using qPCR in NTC vs. KD hSkMC cells (*n = 3* biological replicates). (F) Representative western blot of PRKN protein expression in hSkMC compared to GAPDH control. All graphical and numeric data represent mean ± SEM. \**p<0.05*, \*\**p<0.01*, \*\*\**p<0.001*, \*\*\*\**p<0.0001* vs. NTC by Student’s t-test.

### Overexpression of *TOMM40*, *TOMM22*, and *TOMM20* do not rescue statin-induced decreases in mitochondrial function and content of cholesterol or CoQ

We next sought to determine whether overexpressing *TOMM40* and *TOMM22*, singly and in combination, can rescue simvastatin-induced effects on mitochondrial function. Firstly, we demonstrated that treatment of C2C12 and hSkMC cells with 2 µM simvastatin for 24 hrs resulted in decreased OCR (both basal and maximal oxygen consumption) and ATP production (**Fig. 6A-D)**. Accordingly, mitoROS production was increased by simvastatin in skeletal myotubes (Fig. 6E). Introduction of *TOMM40*- and *TOMM22*-containing lentiviral plasmids, singly and in combination, to these simvastatin-treated cells resulted in no changes in OCR and mitoROS levels (**Fig. 6A-E**). Similarly, over-expressing *Tomm20*, another key component of the TOM complex, did not reverse the simvastatin effect on basal respiration, ATP production, and mitoROS levels in C2C12 cells (**Fig. 6A-E**).

**Figure 6.**
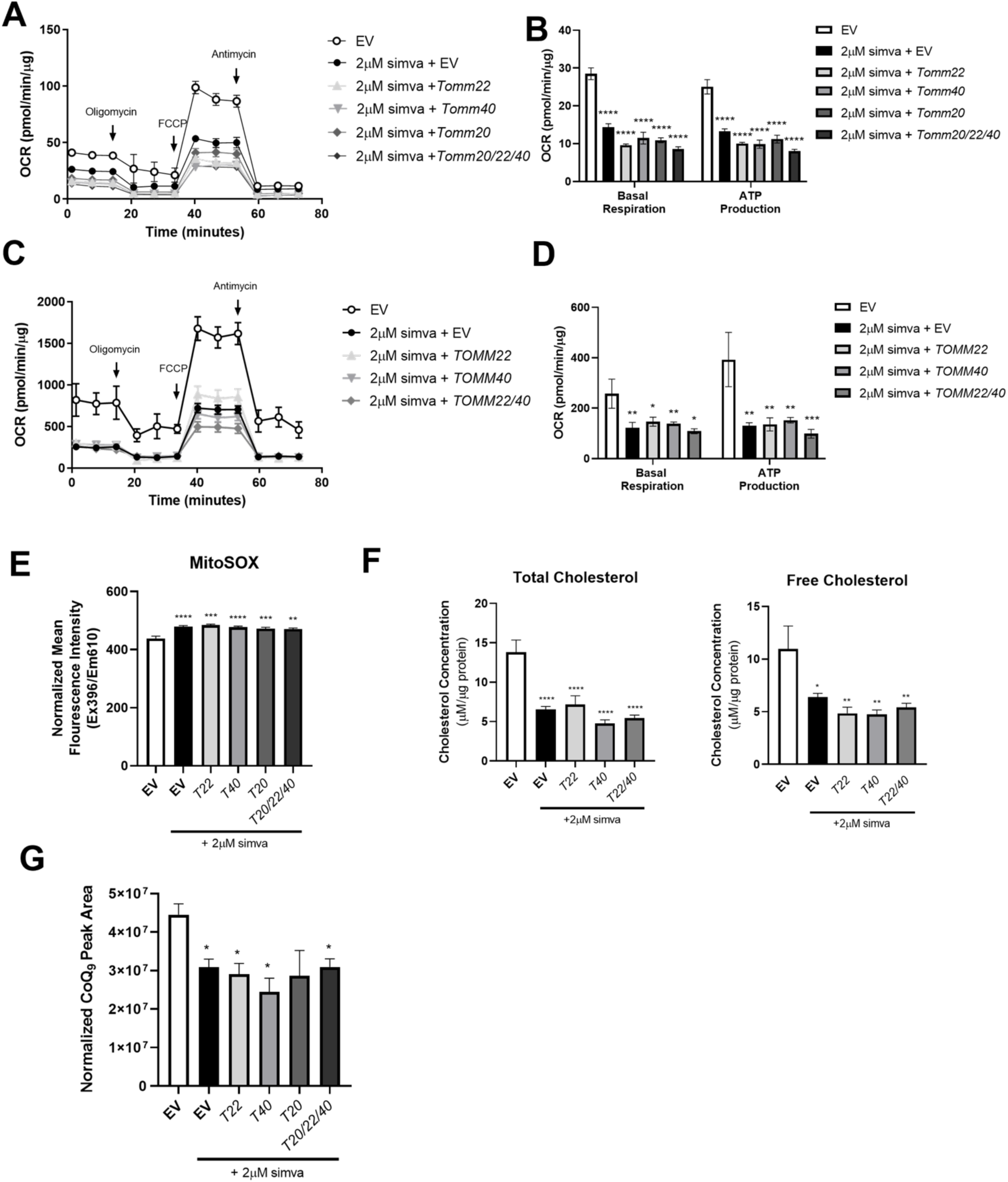
*TOMM40, TOMM22*, and *TOMM20* overexpression, singly and in combination do not alter mitochondrial respiration, ATP production, or MitoSOX, cholesterol or CoQ levels after simvastatin treatment of skeletal muscle cells. (A) OCR was measured in C2C12 cells treated with 2 µM simvastatin + empty vector or *Tomm22*, *Tomm40*, *Tomm20*, or *Tomm20/22/40* expressing plasmids for 24-48 hrs. (B) Basal respiration and ATP production determined from. OCR analysis in (A). (C-D) The same experiment was conducted in hSkMC cells treated with 2 µM simvastatin + empty vector or *TOMM22*, *TOMM40*, *TOMM22/40* expressing plasmids for 24-48 hrs. (*n = 10-12* biological replicate) (E) mitoROS was determined in C2C12 cells by quantifying mean mitoSOX™ fluorescence intensity and normalizing to total protein concentration using Bradford assay (*n = 10-12* biological replicates). (F) Total and free cholesterol were quantified in mitochondria of C2C12 myotubes using Amplex Red Cholesterol Assay (*n = 3* biological replicates). (G) Total CoQ_9_ was quantified from mitochondria isolated from C2C12 cells transfected with empty vector (control), 2 µM simvastatin + empty vector, *Tomm22*, *Tomm40*, *Tomm20*, or *Tomm20/22/40* expressing plasmids (*n =* 3 biological replicates). All values were normalized to protein concentration by BCA. All graphical and numeric data represent mean ± SEM. \**p<0.05*, \*\**p<0.01*, \*\*\**p<0.001*, \*\*\*\**p<0.0001* vs. EV by Welch and Brown-Forsythe or one-way ANOVA, with post-hoc Student’s t-test to identify differences between groups.

In accord with inhibition of mevalonate synthesis by statins^1^, simvastatin (2 µM for 24 hrs) resulted in reduced levels of two products of the mevalonate/CoQ biosynthesis pathway - cholesterol and CoQ - in mitochondria isolated from C2C12 cells by subcellular fractionation (**Fig. 6F, G**). These effects were not reversed by overexpression of *Tomm40* and *Tomm22*, or of *Tomm20.* Together, these results indicate that effects of statin other than downregulation of these genes are primarily responsible for disrupting mitochondrial function and maintaining cholesterol and CoQ content in mitochondria of skeletal muscle cells.

### *TOMM40* and *TOMM22* rescue statin-induced mitochondrial fusion and fission events

As we had observed with *Tomm40* and *Tomm22* KD, TEM image analysis of C2C12 myotubes treated with simvastatin 2 µM for 24 hours (**Fig. 7A**) resulted in reduced mitochondria length and increased width (**Fig. 7B, C**). The likelihood that this resulted from increased mitochondrial fission events was supported by qPCR analysis showing that simvastatin treatment resulted in increased expression of *FIS1* and *DRP1* and reduced expression of *MFN2* and *OPA1* (**Fig. 7D, E**). Furthermore, both mitochondrial density and mtDNA copy number increased in simvastatin-treated C2C12 myotubes compared to control (**Fig. 7F, G**). Thus, as with *TOMM40* and *TOMM22* KD, simvastatin treatment of skeletal myotubes increased mitochondrial number and fragmentation due to increased fission and reduced fusion of these organelles.

**Figure 7.**
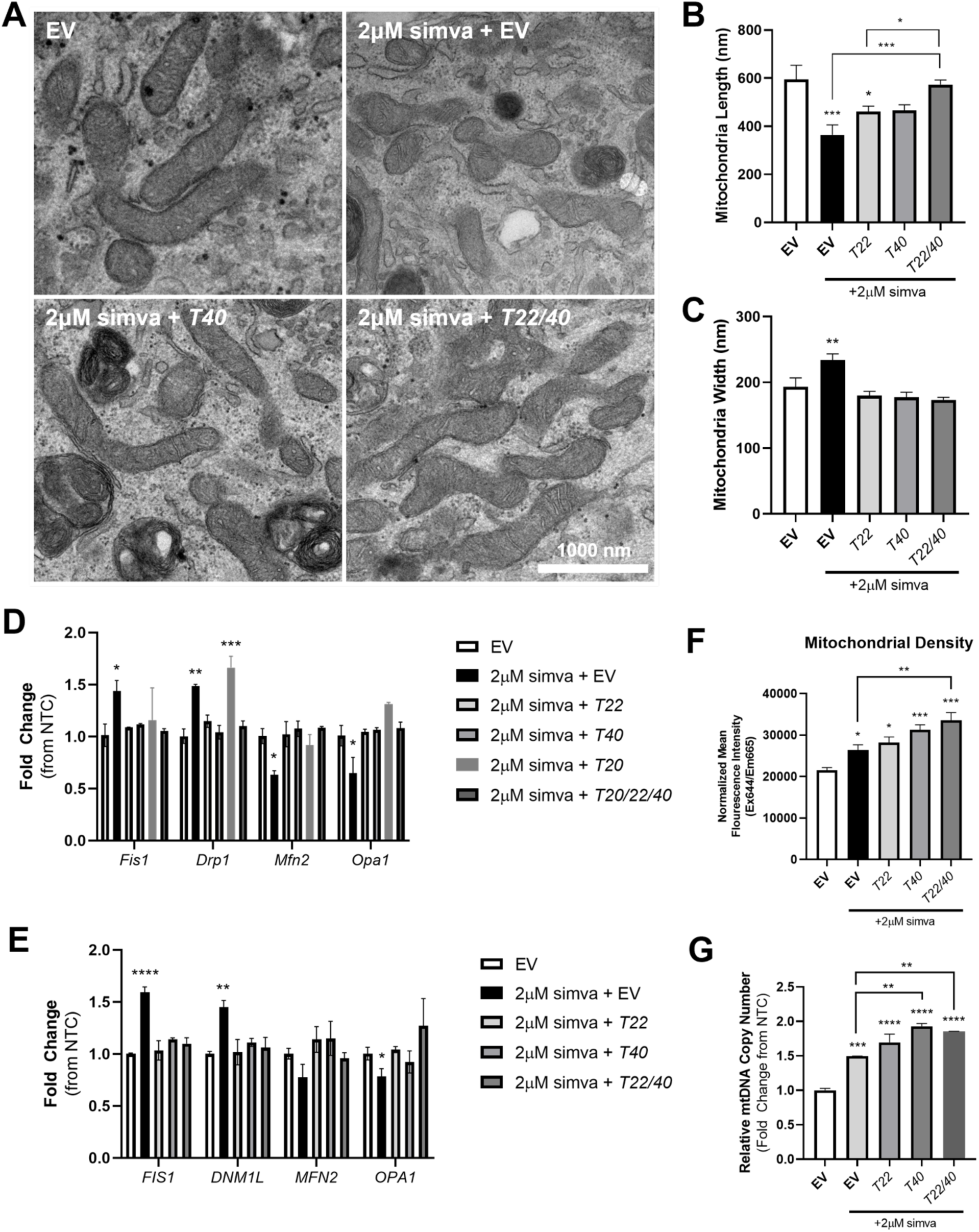
Overexpressing *TOMM40* and *TOMM22, singly and in combination,* suppresses statin-induced mitochondrial fission and promotes fusion in skeletal muscle cells. (A) Representative TEM images of mitochondrial morphology in C2C12 cells transfected with empty vector (EV), 2 µM simvastatin + EV, and *Tomm22 (T22), Tomm40 (T40)*, or *Tomm22/40 (T22/40)* expressing plasmids. (B and C) Using ImageJ software analysis in conjunction with the TEM images in (A), average mitochondrial length and width (nm) were measured (*n = 10-12* cells). (D) Mitochondrial fission (*Fis1/FIS1, Drp1/DNM1L*) and fusion (*Mfn2/MFN2, Opa1/OPA1*) markers were quantified by qPCR in EV and 2 µM simvastatin + EV, *T22, T40*, *Tomm20 (T20)*, and *Tomm20/22/40 (T20/22/40)* overexpressing C2C12 cells. (E) The same experiment was conducted as in (D) but with hSkMC cells. (*n=3* biological replicates) (F) Mitochondrial density was quantified using MitoTracker™ Deep Red FM fluorescence probe to measure average fluorescence intensity and normalized to protein concentration by Bradford assay (n=10-12 biological replicates). (G) Relative mtDNA copy number levels were quantified and normalized to *B2m* transcript levels (nuclear DNA) by qPCR in siRNA transfected C2C12 cells (*n = 3* biological replicates). All graphical and numeric data represent mean ± SEM. \**p<0.05*, \*\**p<0.01*, \*\*\**p<0.001*, \*\*\*\**p<0.0001* vs. EV (without statin) by one-way ANOVA, with post-hoc Student’s t-test to identify differences between groups.

We next tested whether overexpressing *TOMM40* and *TOMM22*, singly and in combination, could rescue these effects of simvastatin on mitochondrial dynamics. Notably, there was a reversal of statin effects on mitochondrial length and width after the addition of *Tomm40* and *Tomm22/40* plasmids in C2C12 myotubes (**Fig. 7B, C**). Moreover, *TOMM40* and *TOMM22* overexpression in both C2C12 and hSkMC cells resulted in reversal of the statin effects on gene expression of the mitochondrial fission and fusion markers described above (**Fig. 7D, E**), while this was not the case for *Tomm20* overexpression in C2C12 myotubes (**Fig. 7D**).

Interestingly, the addback of *Tomm40* and *Tomm22* plasmids to simvastatin-treated cells resulted in further increases in mtDNA copy number and mitochondrial density (**Fig. 7F, G**). Together with the evidence that *TOMM40* and *TOMM22* expression reverse statin effects on mitochondrial dynamics, this suggests that this treatment may suppress statin-induced mitophagy while generating new, healthy mitochondria. Consistent with this hypothesis, we observed by TEM that simvastatin-treated C2C12 cells had a lower percentage of healthy Type 1 mitochondria (sharp cristae and dense matrix) and a higher percentage of Type 2 (dilute cristae and/or dilute matrix) and Type 3 (ruptured) mitochondria, representing abnormal mitochondrial morphology and signs of mitochondrial injury^35^. In addition, there was an increase in the percentage of mitophagosomes per cell in simvastatin-treated skeletal muscle (**Fig. 8D**). Consistent with these observations, we demonstrated increased expression of mitophagy biomarkers *PINK1* and *PARKIN* with simvastatin treatment by qPCR (**Fig. 8E, F**).

**Figure 8.**
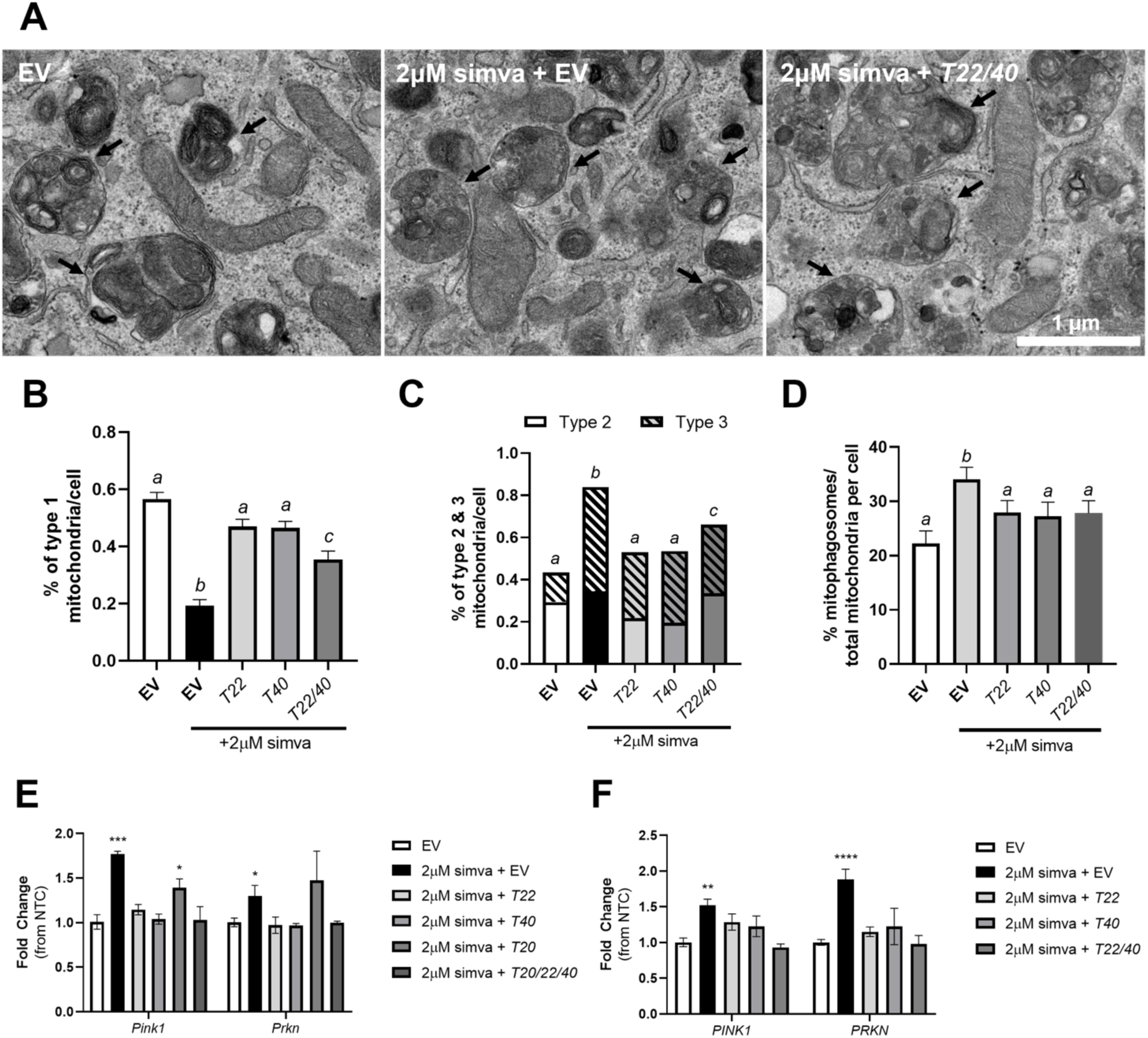
Overexpression of TOMM40 and TOMM22, but not TOMM20, rescues simvastatin-induced mitophagy leading to reduced mitochondrial damage. (A) TEM micrographs of mitophagosomes found in EV, 2 µM simvastatin + EV, and 2 µM simvastatin + *Tomm22/Tomm40* overexpressing C2C12 cells. Arrowheads indicate mitophagosomes. (B-C) Mitochondria in cells were identified and categorized as type 1, 2, or 3 and quantified as a percentage of total mitochondria in each cell. (*n = 10-12* cells). (D) Percent of mitophagosomes per total number of mitochondria per cell, identified from TEM images (*n = 10-12* cells). (E-F) mRNA transcript levels of *Pink1*/*PINK1* and *Prkn/PRKN* (mitophagy), were quantified using qPCR in EV vs. 2 µM simvastatin + EV, *TOMM22, TOMM40, TOMM20*, or *TOMM20/22/40* in (E) C2C12 and (F) hSkMC myotubes (*n = 3* biological replicates). All graphical and numeric data represent mean ± SEM. \**p<0.05*, \*\**p<0.01*, \*\*\**p<0.001*, \*\*\*\**p<0.0001* vs. EV by one-way ANOVA, with post-hoc Student’s t-test to identify differences between groups.

We then showed by TEM that overexpressing *Tomm40* and *Tomm22* in statin-treated C2C12 myotubes resulted in an increase in Type 1 mitochondria and a reduction in Type 2 and Type 3 mitochondria, indicating that *Tomm22* and *Tomm40* are able to rescue, at least in part, simvastatin-induced mitophagy. Consistent with this effect, *TOMM22, TOMM40*, and *TOMM22/40* (but not Tomm20) overexpression reduced *PINK1* and *PARKIN* expression to control levels (**Fig. 8E-F**). In summary, these results demonstrate that simvastatin promotes mitochondrial fission and mitophagy, resulting in an increase in damaged mitochondria, while overexpressing *TOMM22* and *TOMM40* can reverse these effects, thus maintaining mitochondrial quality and homeostasis.

## DISCUSSION

We here report that TOMM40 and TOMM22, key members of the TOM complex, are essential in maintaining mitochondrial function and quality in skeletal muscle cells by promoting their oxidative function, retaining CoQ and cholesterol content, and preserving their morphology and dynamics.

Our finding that mitochondrial function in skeletal myotubes is impaired by suppressing *TOMM40* and *TOMM22* expression is consistent with previous studies in epithelial ovarian cancer and HeLa cells, where knockdown of *TOMM40* disrupted the mitochondrial membrane potential, reduced ATP levels, and elevated mitochondrial ROS levels^36–39^. It has been suggested that mitochondrial dysfunction due to suppression of *TOMM40* and *TOMM22* is caused by interfering with the uptake of mitochondria-targeted proteins^40^. These reports show that approximately 1100 out of 1500 proteins imported by the TOM complex rely on TOMM22^41^. Notably, suppression or mutation of *TOMM22* results in the inactivation of mitochondrial proteins due to misfolding in yeast^41^, as well as apoptosis of human epithelial and endothelial cells, and hepatocytes of zebrafish^42^.

Mitochondria require cholesterol for the maintenance of membrane potential to regulate mitochondrial respiration^32^. We found that *TOMM40* and *TOMM22* KD reduced cholesterol content in skeletal muscle cells and isolated mitochondria, consistent with a report that *TOMM40* KD reduces cholesterol content of HeLa cells^36^, and suggesting that this effect contributes to the mitochondrial dysfunction observed with KD of these TOM components in our study. Interestingly, adding back LDL cholesterol restored mitochondrial cholesterol levels to baseline in *TOMM22* KD skeletal myotubes, but not in *TOMM40* KD cells, suggesting that TOMM40 is necessary for maintaining cholesterol levels in muscle mitochondria^43^. However, neither KD group showed an improvement in mitochondrial OCR after the addition of LDL, indicating that under these conditions, mitochondrial respiration and function are regulated by factors other than cholesterol content, or possibly that the added cholesterol was not introduced into the mitochondrial membrane.

In association with the mevalonate pathway, CoQ_10_ (humans) or CoQ_9_ (in mice) of the CoQ biosynthesis pathway, essential for regulating the mitochondria electron transport chain and thus mitochondrial function, are dependent on isoprenoids, a product of the mevalonate pathway^44^. To date, it is still unclear how CoQ and its precursors are transported across the outer mitochondrial membrane into the matrix of skeletal muscles. However recently, Tai et al.,^45^ showed in *Saccharomyces cerevisiae* that isopentenyl pyrophosphate (IPP) molecules, precursors of both CoQ and cholesterol, may enter the mitochondrial matrix via an IPP transporter situated on the inner mitochondrial membrane. Together with our results showing *TOMM40* and *TOMM22* KD in C2C12 skeletal myotubes resulted in a significant reduction of mitochondrial CoQ content, we suggest TOMM40 and TOMM22 may affect the transport of proteins required for CoQ biosynthesis including those involved in the transport of 4-hydroxybenzoate (4-HB) and isoprenoid pyrophosphates into the mitochondria^46^. Further studies are necessary to determine which transporters and enzymes of the CoQ biosynthesis pathway are recognized and imported by TOMM40 and TOMM22, thus promoting CoQ synthesis in the mitochondria^47^.

Mitochondrial dynamics involves coordination of fusion and fission events that define mitochondria number, size, and morphology^48^. Accumulating evidence reveals a strong association between mitochondrial function and dynamics^49,50^. In addition, genes regulating mitochondrial dynamics, including *MSTO1*, have been shown to impact other cellular functions including oxidative stress, apoptosis, and mitophagy, as well as to induce myopathy and ataxia^51–56^. Since TOMM40 is known to interact with several ER membrane proteins found at mitochondria-ER contact sites (MERCs), including BCAP31 and STAR^56,57^, it is plausible that TOMM40 regulates mitochondrial fusion and fission by directly or indirectly interacting with known markers of fission and fusion localized at MERCs, such as FIS1, DRP1, and MFN2^58^. In support of this, FIS1 has been shown to physically interact with BCAP31 at MERCs, regulating mitochondrial fusion/fission dynamics^59^. Additionally, studies in yeast have shown TOMM22 to interact with PINK1, a key regulator of mitochondrial quality and mitophagy^60–63^. In mammalian cells, dephosphorylation of TOMM22 impairs PINK1 import into the mitochondria, promoting mitophagy^64^.

We show here for the first time, using both TEM and expression of biomarkers, that KD of *TOMM40* and *TOMM22* in skeletal myotubes causes a shift in mitochondrial dynamics towards increased fission events, with a resulting increase in mitophagy as a means of eliminating damaged mitochondria and maintaining mitochondria quality^65,66^. This effect on mitophagy may be mediated, at least in part, by increased mitoROS as a consequence of mitochondrial dysfunction^67,68^. Consistent with this mechanism, we observed that both mitoROS levels and the mitophagy markers *PINK1* and *PRKN* are upregulated with *TOMM40* and *TOMM22* KD.

Based on our evidence for disruption of mitochondrial function and dynamics by *TOMM40* and *TOMM22* knockdown in skeletal muscle cells, and previous evidence that simvastatin downregulates *TOMM40* and *TOMM22* gene expression, we compared the effects of simvastatin exposure with those of *TOMM40* and *TOMM22* knockdown on these properties, and tested whether overexpression of these genes could reverse the statin effects. We showed that simvastatin treatment exerted similar effects on mitochondrial dynamics and morphology in skeletal muscle cells as those observed with both *TOMM40* and *TOMM22* KD. Previous studies in yeast cells reported that statins impair mitochondrial morphology, represented by an increase in aggregated mitochondria, due to disruption of mitochondrial function and membrane potential^69–71^. However, we are the first to show that the reduction in mitochondrial length and increased mitochondrial fragmentation in C2C12 cells exposed to simvastatin can be explained by an increase in mitochondrial fission and decrease in fusion events. As with *TOMM40* and *TOMM22* KD, the shift towards mitochondrial fission resulted in increased mitophagy (e.g. *PINK1* and *PRKN*) to promote the removal of damaged mitochondria^72,73^. Notably, overexpression of *TOMM40* and *TOMM22* rescued the changes in mitochondrial dynamics and morphology, and mitophagy, caused by simvastatin treatment of C2C12 myotubes, suggesting that statin-induced downregulation of these genes may contribute to mitochondrial SAMS. Future work will be required to identify direct and indirect interactions of TOMM40 and TOMM22 with proteins involved in fission, fusion, and mitophagy at MERCs in order to pinpoint key pathways causing the observed changes in mitochondrial dynamics and mitophagy.

Consistent with previous studies^7,9,10^, we showed that simvastatin treatment of C2C12 myotubes resulted in reduction of ATP production and mitochondrial respiration, together with an increase in mitoROS production. We also observed that simvastatin reduced free cholesterol content in whole cells as well as in isolated mitochondria^24^, an effect that might be linked to impaired mitochondrial structure and function. Moreover, as further expected from statin inhibition of the mevalonate pathway, we showed that simvastatin treated C2C12 cells exhibited a reduction in mitochondrial CoQ levels, an effect that has been suggested to contribute to SAMS^74^. These results paralleled those observed with *TOMM40* and *TOMM22* KD, raising the question as to whether reduced expression of these genes contributes to these statin effects in muscle, as was the case for statin-induced changes in mitochondrial morphology and dynamics. However, overexpression of these genes, as well as that of another major TOM complex subunit encoding gene, *TOMM20*^75^, failed to restore statin impairment of mitochondrial CoQ, cholesterol content, and mitochondrial function. In the case of CoQ, the effects of simvastatin and *TOMM40* and *TOMM22* KD were additive (**Fig. S4**). Thus, these effects of statin that may contribute to SAMS are independent of its suppression of expression of genes encoding the TOM complex.

In summary, our results demonstrate that TOMM40 and TOMM22, major members of the TOM complex, have key roles in maintaining mitochondrial function, composition, and dynamics in skeletal muscle cells. They further show that simvastatin treatment of these cells increases mitochondrial fragmentation and that this effect is mediated by suppression of transcription of *TOMM40* and *TOMM22.* However, while there are recent reports that mitochondrial dynamics play a significant role in maintaining mitochondrial function^49–50^, our results show that restoration of dynamics by overexpressing *TOMM40* and *TOMM22* is not sufficient to reverse statin-induced impairment of mitochondrial energy metabolism, indicating concomitant or primary effects of other proposed mechanisms^76–77^. The present study has also shown similar reductions of mitochondrial function and cholesterol and CoQ_10_ content with statin exposure and knockdown of *TOMM40* and *TOMM22,* though reduced expression of these genes does not appear to mediate the statin effects. Finally, our results point to the possibility that altered levels or functions of proteins that interact with the TOM complex at MERCs may predispose to statin induced myopathy.

## ACKNOWLEDGEMENTS

We thank Reena Zalpuri and Danielle Jorgens at the University of California, Berkeley, Electron Microscopy Laboratory for their advice and assistance in electron microscopy sample preparation and data collection. We thank Tommy Tran and Justin Y. Chao for their assistance with the Agilent Seahorse XFe96 Extracellular Flux Analyzer and qPCR.

## SUPPLEMENTAL MATERIAL

**Table S1.**
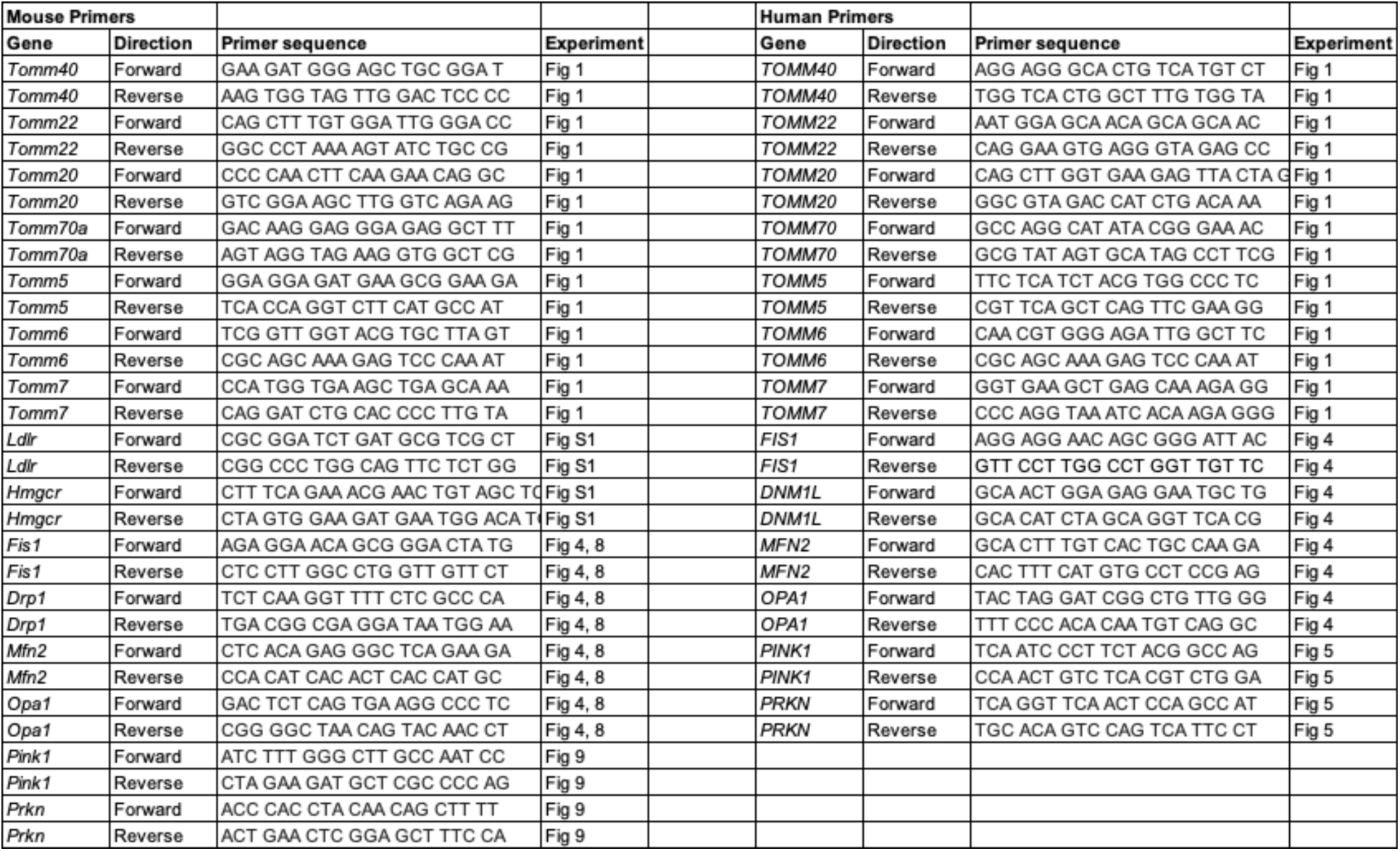
RT-qPCR primers employed in this study.

**Figure S1.**
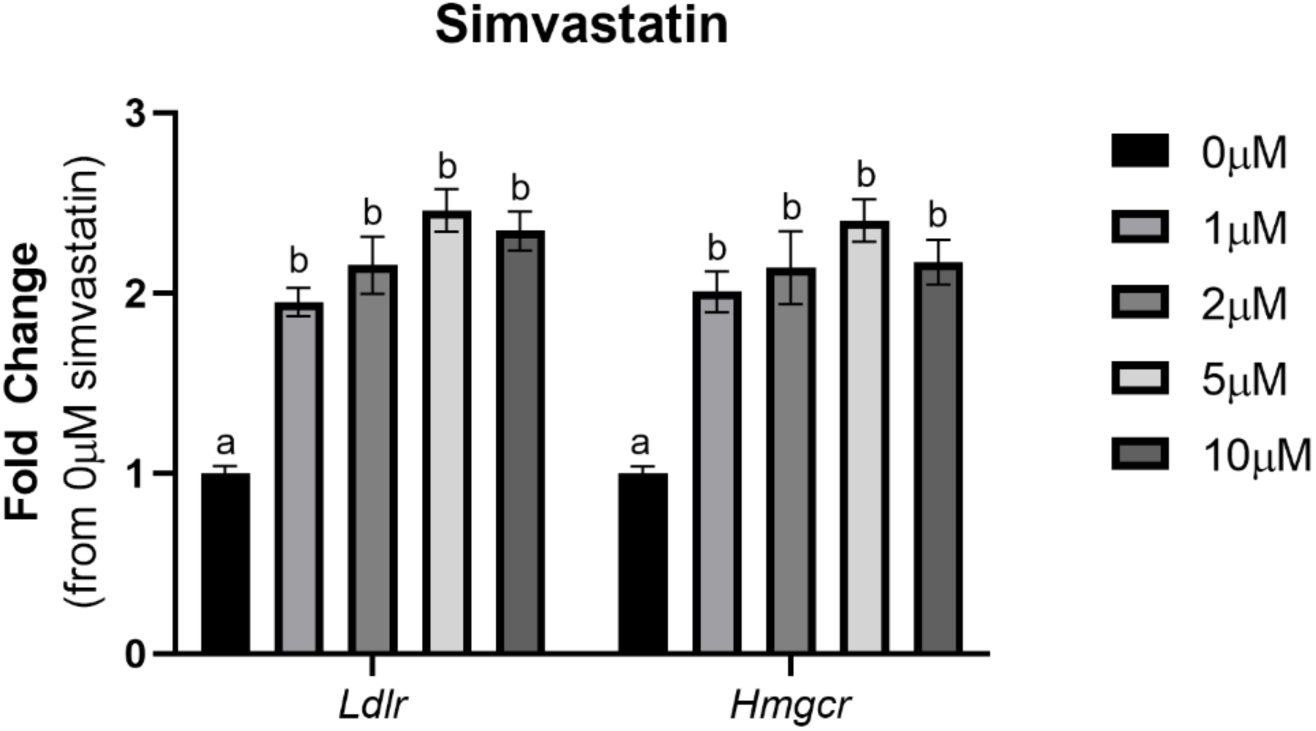
Simvastatin treatment of C2C12 myotubes upregulates *Ldlr* and *Hmgcr* mRNA transcripts in a dose-dependent manner. Differentiated C2C12 myotubes were treated with 0, 1, 2, 5 or 10 µM simvastatin for 48 hrs. All numeric data represent mean ± SEM. \**p<0.05*, \*\**p<0.01*, \*\*\**p<0.001*, \*\*\*\**p<0.0001* vs. NTC by Student’s t-test. (*n = 3* biological replicates)

**Figure S2.**
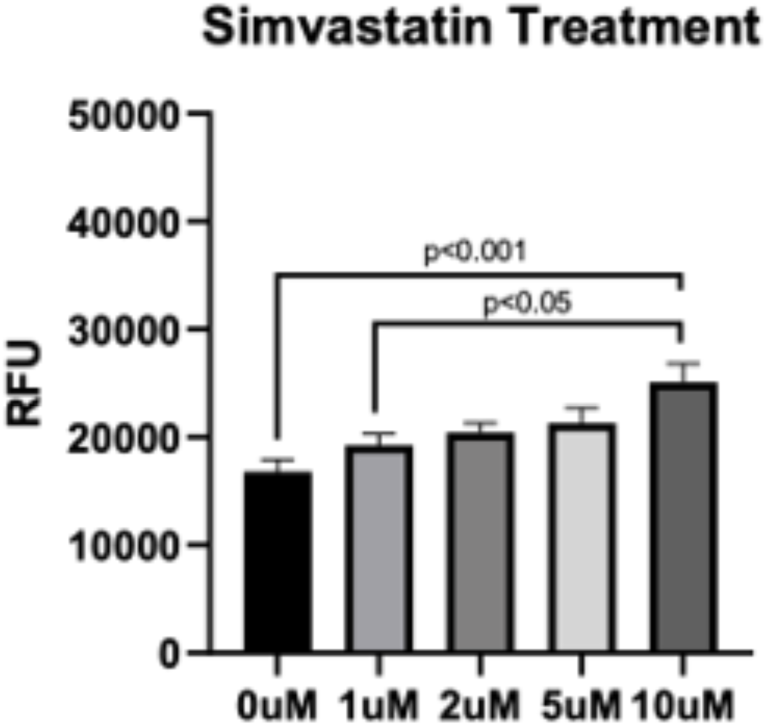
Dose-dependent cell apoptosis identified in C2C12 myotubes. Differentiated C2C12 myotubes were treated with 0, 1, 2, 5 or 10 µM simvastatin for 48 hrs and apoptosis was assessed using an EarlyTox Caspase-3/7 colorimetric detection kit with a spectrophotometer. RFU = relative fluorescence units. All numeric data represent mean ± SEM. \**p<0.05*, \*\**p<0.01*, \*\*\**p<0.001*, \*\*\*\**p<0.0001* vs. NTC by Student’s t-test. (*n = 3* biological replicates)

**Figure S3.**
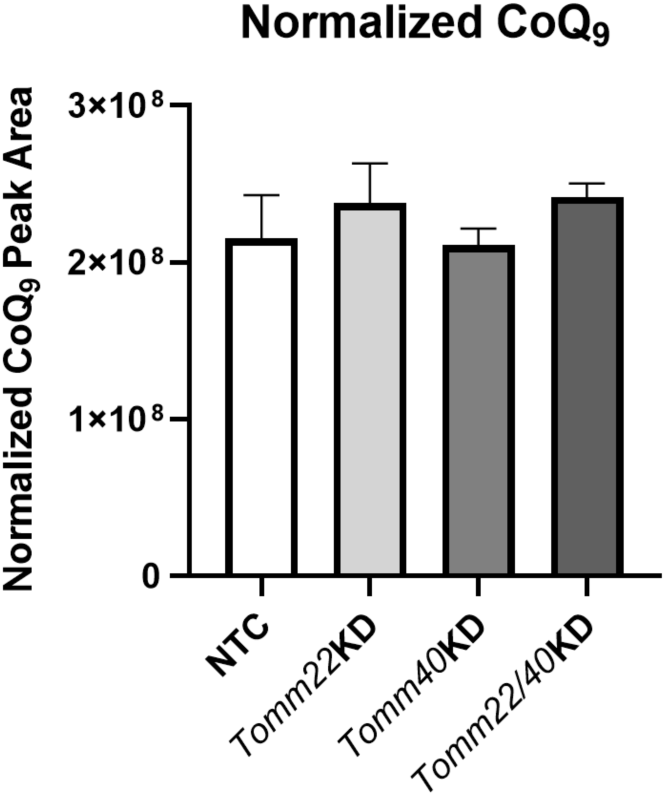
Effects of *Tomm40* and *Tomm22* knockdown on CoQ levels in C2C12 whole cell lysates. CoQ9 from whole cell lysates of NTC, *Tomm40, Tomm22*, and *Tomm22/40* KD C2C12 skeletal myotubes. All values were normalized to protein concentration. All numeric data represent mean ± SEM. \**p<0.05*, \*\**p<0.01*, \*\*\**p<0.001*, \*\*\*\**p<0.0001* vs. NTC by Welch and Brown-Forsythe ANOVA. (*n = 3* biological replicates)

**Figure S4.**
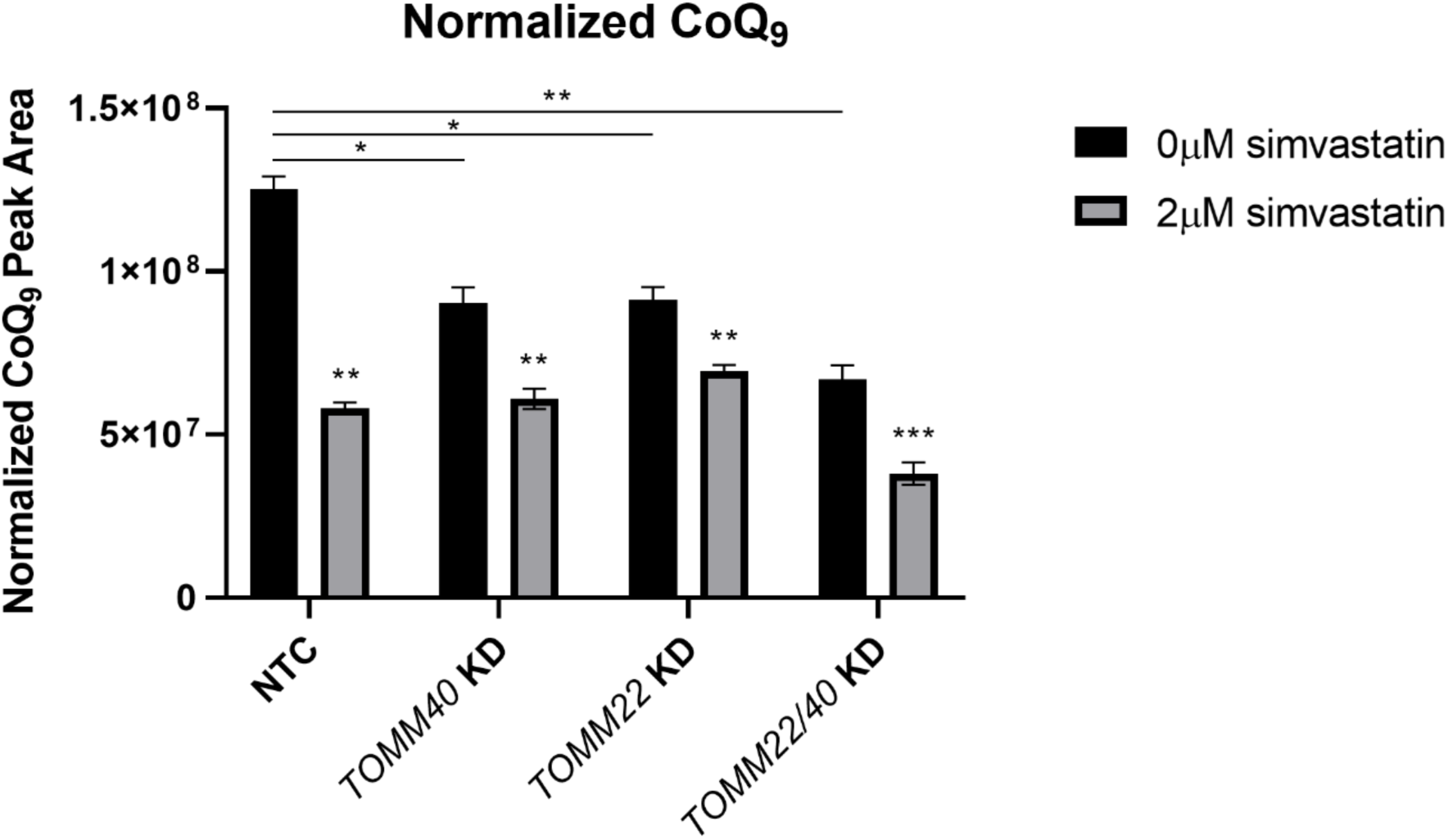
Simvastatin treatment further decreases CoQ levels in *Tomm40* and *Tomm22* KD C2C12 mitochondria. (A)Total CoQ_9_ from isolated mitochondria of NTC, *Tomm40, Tomm22*, and *Tomm22/40* KD C2C12 skeletal myotubes treated with simvastatin (2 µM). All values were normalized to protein concentration. All numeric data represent mean ± SEM. \**p<0.05*, \*\**p<0.01*, \*\*\**p<0.001*, \*\*\*\**p<0.0001* vs. NTC by Welch and Brown-Forsythe ANOVA. (*n = 3* biological replicates)

## NOVELTY AND SIGNIFICANCE

### What is Known?

- Statins disrupt mitochondrial function resulting in increased reactive oxygen species, increased mtDNA fragmentation, inhibition of CoQ_10_ synthesis, and impaired electron transport chain, however the mechanisms for these effects are unclear.
- Simvastatin exposure of skeletal myotubes downregulates genes encoding mitochondrial proteins, including two major subunits of the TOM complex, *TOMM40* and *TOMM22*.

### What New Information Does This Article Contribute?

- Knockdown (KD) of *TOMM40* and *TOMM22* in skeletal myotubes displayed mitochondrial phenotypes similar to those resulting from simvastatin treatment, including decreased oxidative phosphorylation, reactive oxygen species, and mtDNA fragmentation, and decreased mitochondrial CoQ and cholesterol content.
- Both simvastatin treatment and *TOMM40* and *TOMM22* KD disrupted mitochondrial dynamics by promoting mitochondrial fission, reducing fusion, and upregulating mitophagy in response to mitochondrial damage.
- Overexpressing *TOMM40* and *TOMM22* rescued statin-induced effects on mitochondrial fission and fusion, leading to reduction in mitophagy.

Statins are the most commonly used drugs for lowering plasma LDL cholesterol and cardiovascular disease risk. While highly effective and generally well-tolerated, statins can induce clinically significant myopathy, an effect that has been attributed in part to impairment of mitochondrial function, although the underlying mechanism remains unclear. Here we studied two genes encoding major components of the translocase of the outer mitochondrial membrane (TOM) complex, *TOMM40* and *TOMM22*, which were downregulated by simvastatin in C2C12 and hSkMC skeletal myotubes. We demonstrated that knockdown (KD) of these genes in myotubes impaired mitochondrial function, increased mitochondrial superoxide production, and reduced mitochondrial cholesterol and CoQ levels, effects that were also observed with simvastatin treatment. Furthermore, transmission electron microscopy showed that both *TOMM40* and *TOMM22* KD and simvastatin exposure of skeletal myotubes disrupted mitochondrial dynamics and morphology, consistent with expression of biomarkers of increased mitochondrial fission and decreased fusion, and evidence for a secondary increase of mitophagy. Although overexpressing *TOMM40* and *TOMM22* did not rescue simvastatin effects on mitochondrial function or cholesterol and CoQ content, there was rescue of statin effects on mitochondrial dynamics, morphology, and mitophagy. These findings are consistent with TOMM40 and TOMM22 as key regulators of mitochondrial homeostasis and suggest that their downregulation by statins may contribute to myopathy by disrupting mitochondrial dynamics.

